# Structural basis for excitatory neuropeptide signaling

**DOI:** 10.1101/2023.04.29.538817

**Authors:** Valeria Kalienkova, Mowgli Dandamudi, Cristina Paulino, Timothy Lynagh

**Author notes:** Correspondence and requests for materials should be addressed to CP, or TL,. Department of Biomedicine, University of Bergen, Bergen, Norway (Valeria Kalienkova). Heidelberg University Biochemistry Center (BZH), Im Neuenheimer Feld 328, 69120 Heidelberg, Germany.

## Abstract

Rapid chemo-electric signaling between neurons is mediated by ligand-gated ion channels, cell-surface proteins with an extracellular ligand-binding domain and a membrane-spanning ion channel domain^1^. The degenerin/epithelial sodium channel (DEG/ENaC) superfamily, which occurs throughout the animal kingdom, is unique in its diversity of gating stimuli, with some DEG/ENaCs gated by conventional ligands such as neuropeptides, and others gated by e.g. pH, mechanical force, or enzymatic activity^2-5^. The mechanism by which ligands bind to and activate DEG/ENaCs is poorly understood. We have therefore dissected the structural basis for neuropeptide binding and gating in a neuropeptide-gated DEG/ENaC, FMRFamide-gated sodium channel 1 (FaNaC1) from the annelid worm *Malacoceros fuliginosus*^6^, using cryo-electron microscopy. High-resolution structures of FaNaC1 in the ligand-free resting state and in several ligand-bound states reveal the ligand-binding site and capture the ligand-induced conformational changes that mediate channel gating. Complementary mutagenesis experiments confirm the functional roles of particular amino acid residues implicated by the structures. Our results illuminate ligand-induced channel gating in DEG/ENaCs and offer a structural template for the experimental dissection of channel pharmacology and ion conduction in a characteristically metazoan ion channel superfamily.

## INTRODUCTION

Ligand-gated ion channels (LGICs) are cell membrane proteins that convert extracellular chemical signals into transmembrane ionic current, thus contributing to rapid inter-cellular signaling and chemo-sensation^1,2^. The major superfamilies of pentameric LGICs, e.g. GABA_A_ receptors and nicotinic receptors, and tetrameric ionotropic glutamate receptors, e.g. AMPA receptors and NMDA receptors, are found in prokaryotes and eukaryotes, and most of these channels are gated by small amino acid or biogenic amine ligands^1,7^. This contrasts with a third LGIC superfamily that is more specific to animals and close relatives, the trimeric degenerin/epithelial sodium channels (DEG/ENaCs)^2-5,8^. Despite having arisen relatively recently, DEG/ENaCs are especially diverse in terms of stimuli, as the superfamily includes constitutively active channels, pH-gated channels, osmolarity-gated channels, mechanically gated channels, and neuropeptide-gated channels, among others^2,3^. DEG/ENaCs are most highly expressed in neurons, where their gating causes depolarization due to selective cation permeability, with moderate selectivity for Na^+^ over K^+^ ions^9,10^.

The fact that such diverse stimuli activate DEG/ENaCs raises several questions, ranging from evolutionary, to physiological, to biophysical. For example, did sensitivity to different ligands emerge independently and on demand in different animal lineages? And from a biophysical perspective, is there a gating machinery unique to the DEG/ENaC architecture that converts very different biophysical stimuli into similar conformational change at the channel gate? So far our knowledge of DEG/ENaC channel architecture and gating derives mostly from X-ray or cryo-electron microscopy (cryo-EM) data^11,12^ and complementary biophysical experiments^13^ on vertebrate acid-sensing ion channels (ASICs), a family of proton-gated DEG/ENaCs. These, together with recent structures of the ENaC extracellular domain, show that DEG/ENaCs are assembled by three homologous subunits, each with a channel-forming transmembrane domain and a large extracellular domain, in threefold symmetry around a central pore^14,15^. As inferred from high-resolution structures of chicken ASIC1, channel gating involves the following conformational changes in each subunit. The protonation of numerous side chains leads to the collapse of a large part of the extracellular domain, whereby the mid-peripheral domain (“thumb”) is drawn upwards towards the outer “finger” domain; concomitantly, β-strands of the low-peripheral “palm” and “wrist” domains move outwards, pulling channel-forming α-helices peripherally to open the channel^12,14^.

It is unknown whether protons and more tangible transmitters, like neuropeptides, induce the same biophysical mechanism of channel gating in cognate DEG/ENaCs. The relationship between neuropeptide-gated and other DEG/ENaCs is also interesting from an evolutionary perspective, as neuropeptide-based, paracrine signaling systems potentially pre-dated and gave rise to more modern synaptic systems^16,17^, and neuropeptide-gated DEG/ENaCs occur in distinct animal types that diverged a long time ago^18,19^. This raises the possibility that neuropeptide-gated channels constitute one of the earliest occurring DEG/ENaCs and that understanding their ligand-induced gating may unfurl broad insights into mechanisms of DEG/ENaC function. Two distinct families of neuropeptide-gated DEG/ENaCs have been described so far. These include, from radially symmetric hydrozoans, the hetero-trimeric pyroQWLGGRFamide-gated Na^+^ channels (HyNaCs)^20^, and from bilaterally symmetric molluscs and annelids, the homo-trimeric FMRFamide-gated Na^+^ channels (FaNaCs)^6,19^. The short neuropeptide FMRFamide (H-Phe-Met-Arg-Phe-NH_2_, “FMRFa”) is of particular significance in bilaterian animals, as its broad neural expression makes it a common marker of the nervous system in model invertebrates such as molluscs, annelids, arthropods and nematodes, in which it mediates signaling via FaNaCs and/or G-protein-coupled receptors^21^.

To uncover the structural basis for excitatory neuropeptide activity and establish principles of ligand recognition and channel gating in the DEG/ENaC superfamily, we have investigated the structure of FaNaC1, an FMRFa-gated DEG/ENaC from the annelid *Malacoceros fuliginosus*, using cryo-EM. We solved high-resolution structures of FaNaC1 alone, with full agonist FMRFa, with partial agonist ASSFVRIa, and with both FMRFa and pore-blocker diminazene, identifying the ligand-binding site and elucidating the conformational changes induced by ligand-binding. Together with complementary mutagenesis and electrophysiological experiments, these results establish ligand recognition and channel gating mechanisms and offer a structural template for the experimental dissection of function throughout the DEG/ENaC channel superfamily.

## RESULTS

### FaNaC structural architecture

To investigate the structure of neuropeptide-gated DEG/ENaC channels, we sought a prototype showing typical function and high heterologous expression. We therefore transfected HEK293T cells with two previously characterized FaNaCs from different branches of the FaNaC family, *Octopus bimaculoides* FaNaC from the mollusc-specific branch, and *Malacoceros fuliginosus* FaNaC1 from the annelid-specific branch^6^, and probed expression via a C-terminal fluorescent tag. With greater expression (not shown) and typical FMRFa-gated Na^+^-selective currents (Fig. 1a), we pursued the structural characterization of *Malacoceros* FaNaC1. We transduced HEK-293S cells, purified FaNaC1, and incorporated into lipid nanodiscs for subsequent cryo-EM study. Preparations of FaNaC1 alone and with FMRFa (30 μM) yielded 3D reconstructions with a global resolution of 2.7 and 2.5 Å, respectively (Fig. 1b,c, Table S1, Fig. S1,S2). For both structures, density could be unambiguously assigned to amino acid sequence based on mostly continuous main chain density and numerous distinctive side chain densities (Fig. S1,S2). The 63-amino acid C-terminal tail was not resolved, consistent with most of this domain being flexible, dispensable, and highly variable across DEG/ENaCs (e.g. ^14,22^).

**Figure 1.**
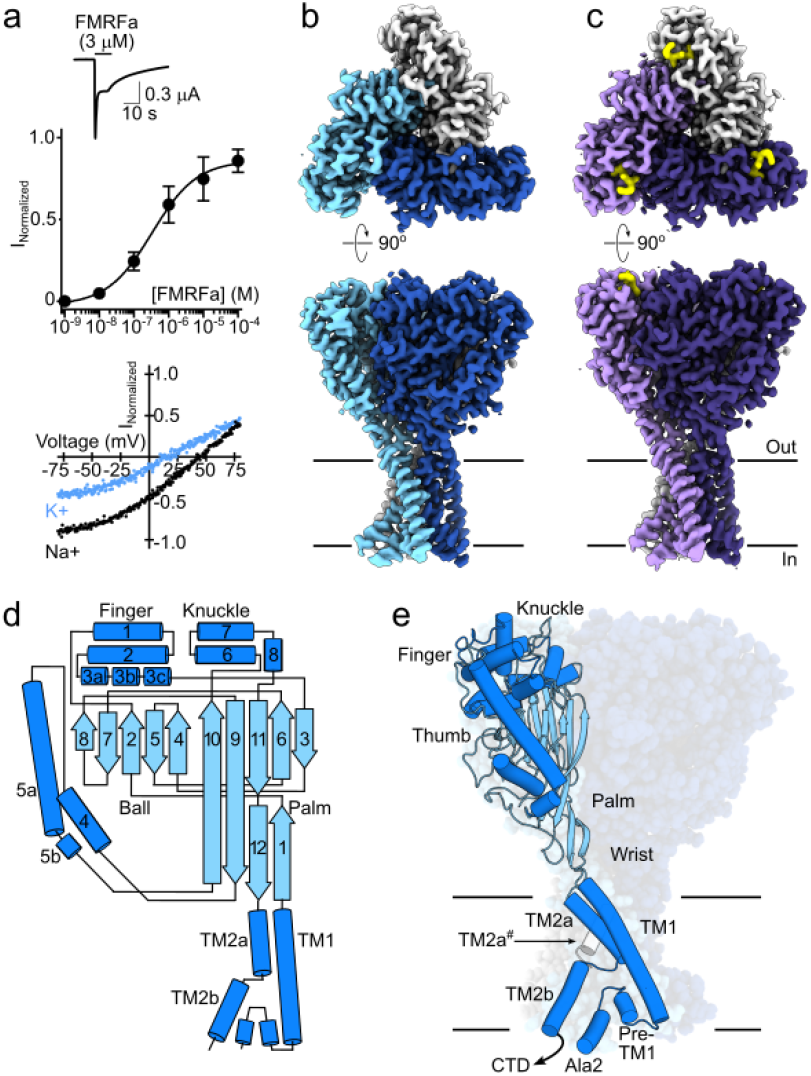
Malacoceros fuliginosus FaNaC1 architecture. (a) FMRFa-gated current (upper), FMRFa concentration-current relationship (mean ± SEM, n = 4, mid), and FMRFa-gated current-voltage relationship in extracellular Na^+^ or K^+^ (lower) in *Xenopus* oocytes expressing FaNaC1. (b,c) Cryo-EM density maps of FaNaC1 (subunits in blues) and FaNaC1/FMRFa (subunits in purples, FMRFa in yellow), viewed from the extracellular side (upper panels) and from within the lipid bilayer (lower panels). Lines indicate bilayer. (d) Schematic of major secondary structure elements. α-helices blue, β-strands cyan. (e) FaNaC1 model highlighting one subunit, colored as in d. *Ala2*, amino acid residue immediately following starting methionine. *CTD*, C-terminal tail, not resolved. TM2a# : adjacent subunit.

The structures confirm that FaNaCs are trimeric like other DEG/ENaCs^14,15^, with three subunits forming central channel pore (Fig. 1b,c). Each subunit comprises a minimal N-terminal and a non-resolved 63-amino acid C-terminal facing the intracellular side, two transmembrane segments (TM1 and TM2), and a large extracellular domain (Fig. 1d,e). The extracellular domain can be divided into palm, thumb, finger, and knuckle domains similar to those previously described for ASIC^14^ and ENaC^15^ (Fig. 1e). TM2 is unwound at a GIS motif in the middle of the membrane, yielding discontinuous upper “TM2a” and lower “TM2b” α-helical segments within each subunit (Fig. 1e, Fig. S1,S2). Consequently, TM2a from one subunit essentially forms a membrane-spanning helix with TM2b from the adjacent subunit (Fig. 1e). The upper, middle, and lower segments of the channel pore are lined by TM2a, the GIS motif, and a re-entrant loop from the short pre-TM1 N-terminal segment, respectively (Fig. 1e). Thus, the channel architecture of FaNaC1 is similar to that of its distant DEG/ENaC cousin, ASIC1^11,23^, suggesting that this architecture is probably adopted by most channels of the DEG/ENaC superfamily. We also observe that the hydrophobic periphery of the channel is thinner than the membrane bilayer, such that the outer leaflet of the membrane bends to make way for hydrophilic, lateral fenestrations between adjacent subunits, creating a path for water and ions into the channel pore (Fig. S3).

### Ligand-binding site

In the FaNaC1/FMRFa structure, we observed a discrete cryo-EM density in a small pocket at the upper corner of each subunit, which fits a single FMRFa molecule (Fig. 2a, Fig. S2). The FMRFa-binding pocket is formed by α-helical segments α1 – α3a (residues V87 to F144), the β6-β7 loop of the same subunit (residues D234 to G241), and partly by α6 from the adjacent subunit (G423-K428, Fig. 2a). This is strikingly different from the site we previously proposed based on mutagenesis of highly conserved residues at the mid-extracellular interface of adjacent subunits^6^. The basis for ligand recognition appears to be mostly hydrophobic interactions. The FMRFa N-terminal phenylalanine residue (F1) is positioned atop the pocket and the M2 side chain orients downward between α2-F129, β6-β7 loop-I236 and M238, and α6-F431. FMRFa R3 orients upwards: the density for the guanidino moiety in our map is relatively weak, but the modeled side chain is 4-6 Å from polar side chains α1-D101 and β6-β7 loop-E235 and R237. Finally, FMRFa F4 and C-terminal amide sit deep in the pocket, with the F4 side chain surrounded as closely as 3.5-4.6 Å by the hydrophobic side chains of α1-F97, P103, α2-V122, A126, and F129, and β6-β7 loop-P240. Additionally, FMRFa F1 and M2 main chain carbonyl oxygens are 2.5-3.1 Å from the FaNaC1 α2-Q133 amide side chain (Fig. 2a), indicative of a potential polar interaction.

**Figure 2.**
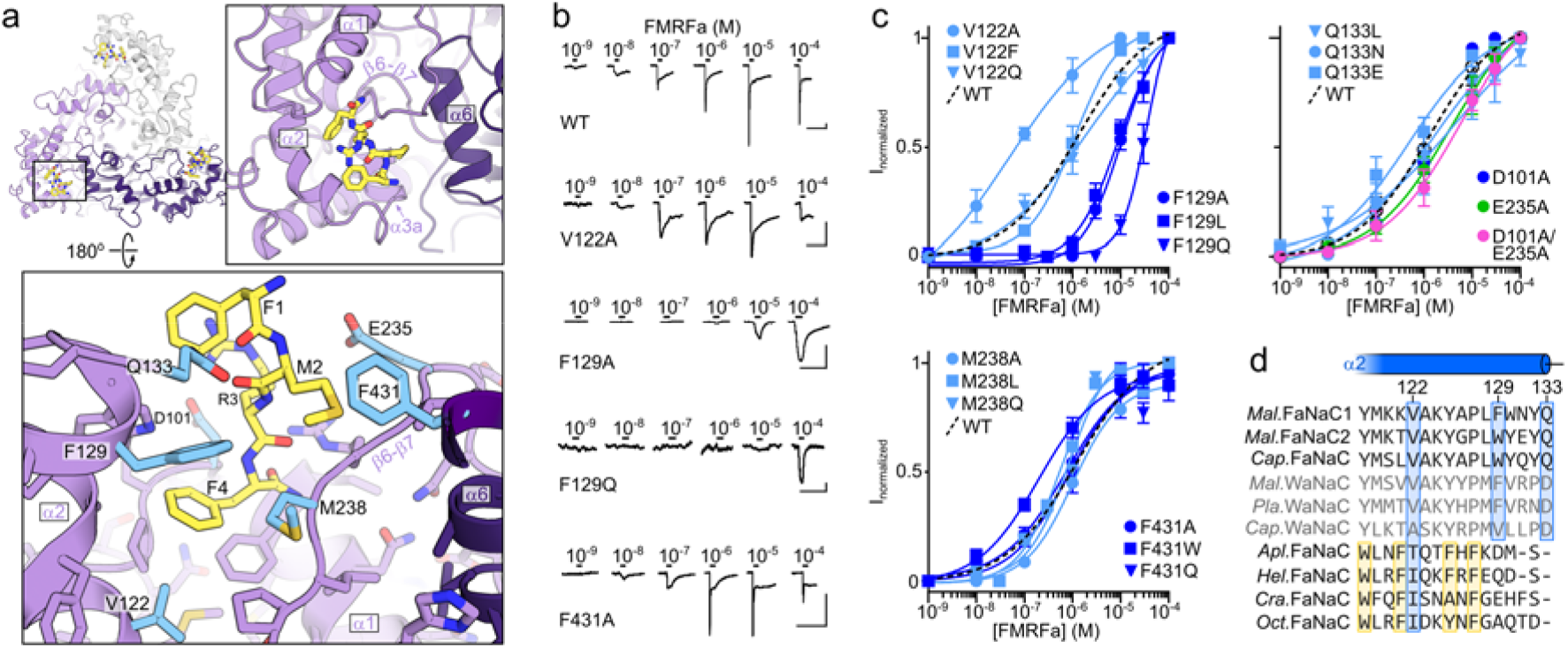
FMRFa binding site. (a) Ligand-binding site in FaNaC1/FMRFa structure. Subunits shaded in purples, FMRFa yellow. Residues tested via mutagenesis cyan and labelled. (b) Example FMRFa-gated currents in oocytes expressing indicated wildtype (WT) and mutant channels. Scale bars: x, 30 s; y, 1 μA (WT, V122A, F129A) or 200 nA (F129Q, F431A). (c) Mean (± SEM, n = 4) normalized currents in response to increasing FMRFa concentrations. (d) Sequence alignment of outer α2-helical segment, including previously characterized FaNaCs (black) and Wamide-gated Na^+^ channels (WaNaCs, grey) from annelids *Mal*., *Malacoceros fuliginosus*; *Cap*., *Capitella teleta; and Pla*., *Platynereis dumerilii;* and molluscs *Apl*., *Aplysia kurodai*; *Hel*., *Helix aspersa*.; *Cra*., *Crassostrea gigas*.; *Oct*., *Octopus bimaculoides*. Blue boxes highlight relative conservation of FaNaC1 ligand-binding residues, yellow boxes residues whose mutation in *Apl*. FaNaC decrease FMRFa potency^24^.

We questioned the divergence of neuropeptide-gated FaNaCs from other DEG/ENaCs by examining this pocket in high-resolution structures of vertebrate ENaC and ASIC. Although α6 lies in a similar position in each channel, α1-α3 arrangement is vastly different in FaNaC1, ENaC, and ASIC (Fig. S4). Consequently, the FMRFa site is essentially obscured by an α-helix in ENaC and by a loop in ASIC (Fig. S4). This suggests that the FMRFa-binding pocket is unique to the FaNaC family, and the enhancement of proton-gated currents that FMRFa elicits in ASICs^39^ likely derives from binding to elsewhere on the channel, consistent with the central, extracellular vestibule peptide-binding site proposed by others for ASICs^40,41^.

To establish the functional importance of interactions apparent in our FaNaC1/FMRFa structure, we compared FMRFa potency at wildtype (WT) and 18 mutant channels with amino acid substitutions at selected positions, via heterologous expression in Xenopus laevis oocytes and two electrode voltage clamp (Fig. 2b,c). Mutations in the α2 helix had the largest effects on FMRFa potency, with F129L and -A mutations decreasing potency 10- to 20-fold (EC_50_: WT, 850 ± 270 nM; F129L, 9 ± 3 μM, F129A, 18 ± 8 μM; each n = 4) and F129Q decreasing potency ∼100-fold (Fig. 2b,c), suggesting that van der Waals interactions between FaNaC1 F129 and FMRFa M2 and F4 contribute substantially to FMRFa binding. Whereas V122F and -Q mutations had no effect on FMRFa potency, V122A caused a 20-fold increase in FMRFa potency (Fig. 2b,c; EC_50_ 40 ± 10 nM, n = 4). The putative polar interactions we probed via mutagenesis make, at most, relatively subtle contributions to FMRFa binding. For example, Q133N and -L mutations, essentially retracting or removing a hydrogen bond partner from the FMRFa main chain, and D101A/E235A, removing two oppositely charged binding partners of the FMRFa R3 side chain, caused two-to four-fold decreases in potency (Fig. 2c). Finally, on the inner wall of the binding pocket, M238 and F431 mutations also had little if any effect on FMRFa potency (Fig. 2c). Thus, FMRFa binding seems to rely mostly on hydrophobic interactions with side chains of the α2 helix.

### Partial agonists bind via a similar mechanism to FMRFa

In addition to FMRFa, several other products of neuropeptide precursors can gate certain FaNaCs. For example, FVRIamides gate annelid FaNaCs, and FLRFa gates multiple mollusc FaNaCs, but generally with lower potency and efficacy than FMRFa^6,19,25,26^. We examined the structural basis of this partial agonism by solving the cryo-EM structure of FaNaC1 in the presence of ASSFVRIa, a product of the FVRIamide precursor in several annelids that gates FaNaC1 with relatively low potency and efficacy (Fig. 3a). The FaNaC1/ASSFVRIa cryo-EM map was resolved to 2.4 Å resolution, with a discrete density in the same ligand-binding pocket as that described for FMRFa (Fig. 3b; Fig. S5). This density was best fit by the C-terminal FVRIa segment of the peptide, and the N-terminal ASS segment was not resolved (Fig. 3c,d, Fig. S5). This indicates a very similar binding mechanism for both full and partial agonists, whereby FaNaC1 V122 and F129 coordinate the hydrophobic C-terminal side chain— F4 in FMRFa and I7 in ASSFVRIa (Fig. 2a and Fig. 3c,d). We confirmed that the four C-terminal residues of ASSFVRIa determine its agonist activity by measuring FaNaC1 responses to FVRIa, observing very similar activity to the parent peptide (Fig. 3a). Similar binding mechanisms for neuropeptides with loosely conserved hydrophobic C-terminal residues and divergent N-terminal segments explains how annelid FaNaCs are gated by diverse neuropeptides, including FMRFa, various FVRIamides, and LFRYa^6^.

**Figure 3.**
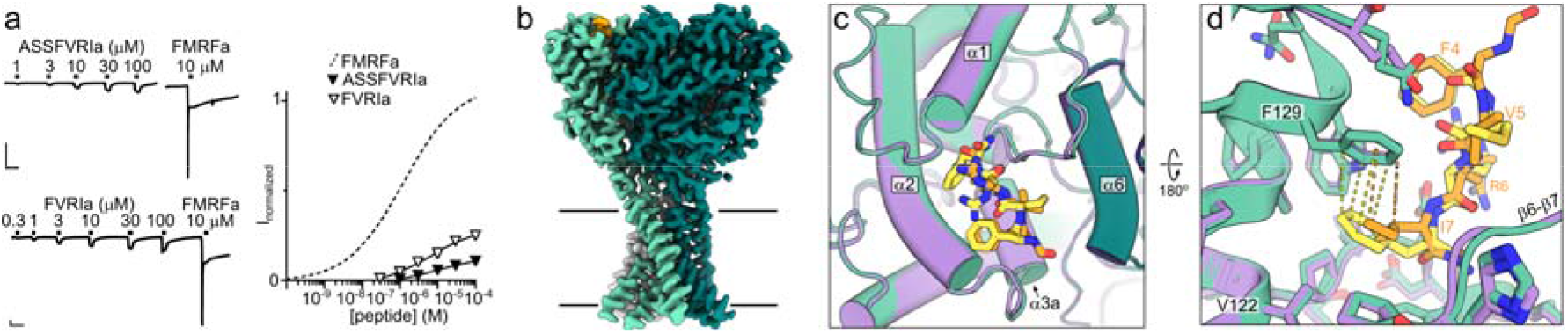
Partial agonist binding in the FMRFa site. (a) *Left*, FaNaC1 currents gated by ASSFVRIa, FVRIa, and FMRFa, and *right*, average (± SEM, n = 4 or 5) ASSFVRIa- and FVRI-gated current amplitude normalized to FMRFa. Scale bars: x, 30 s; y, 250 nA. (b) Cryo-EM map of FaNaC1/ASSFVRIa. FaNaC1 subunits shaded in greens, ASSFVRIa yellow. (c) Overlay of FaNaC1/FMRFa (purple) and FaNaC1/ASSFVRIa (teal) models viewed from above. (d) Magnified view of overlay from (c) viewed from below. Dashed lines indicate 3.3-4.5 Å FaNaC1 F129—FMRFa F4 (yellow) and FaNaC1 F129—ASSFVRI I7 (orange) inter-atom distances.

Full agonist FMRFa and partial agonist ASSFVRIa induce highly similar FaNaC1 conformations (Fig. 3c). In addition to the high similarity between FaNaC1/FMRFa and FaNaC1/ASSFVRIa structures, neither preparation yielded additional distinct 3D classes or any indication of conformational heterogeneity in the image processing (Fig. S2, S5). Thus, the partial agonism of ASSFVRIa does not appear to derive from the induction of a different conformational change compared to FMRFa. Instead ASSFVRIa might have lower affinity than FMRFa especially for the active channel state, which could cause decreased gating efficacy^27^. Indeed, ASSFVRI I7 lacks energetically favourable π-stacking interactions available to FMRFa F4, in addition to fewer individual hydrophobic interactions (Fig. 3d). Furthermore, the density for ASSFVRIa F4 is less definitive than that of the equivalent FMRFa F1 residue despite higher overall resolution of the FaNaC1/ASSFVRIa map (Fig. S2, S5), further indicative of ASSFVRIa binding less tightly.

### A putative open-channel state

To establish how ligand-binding induces channel gating, we compared ligand-free and ligand-bound FaNaC1 structures from the binding pocket down to the channel pore. In both ligand-bound structures, however, the channel pore appears closed, with radii of ∼1 Å at the level of G503 and G506 in TM2a (G3’ and G6’ in a TM2 numbering scheme^28^; Fig. 4a). This is similar to our ligand-free, inactive FaNaC1 structure (Fig. 4a) and is too narrow to pass even mostly dehydrated Na^+^ ions (∼2.3 Å). Thus, FMRFa- and ASSFVRIa-bound channels have probably adopted a desensitized state in the prolonged presence of agonist, in accord with the large decrease in current amplitude that occurs within ∼2 s of FMRFa application, especially at higher FMRFa concentrations (e.g. Fig. 1a, 2b, 3a, 4b).

**Figure 4.**
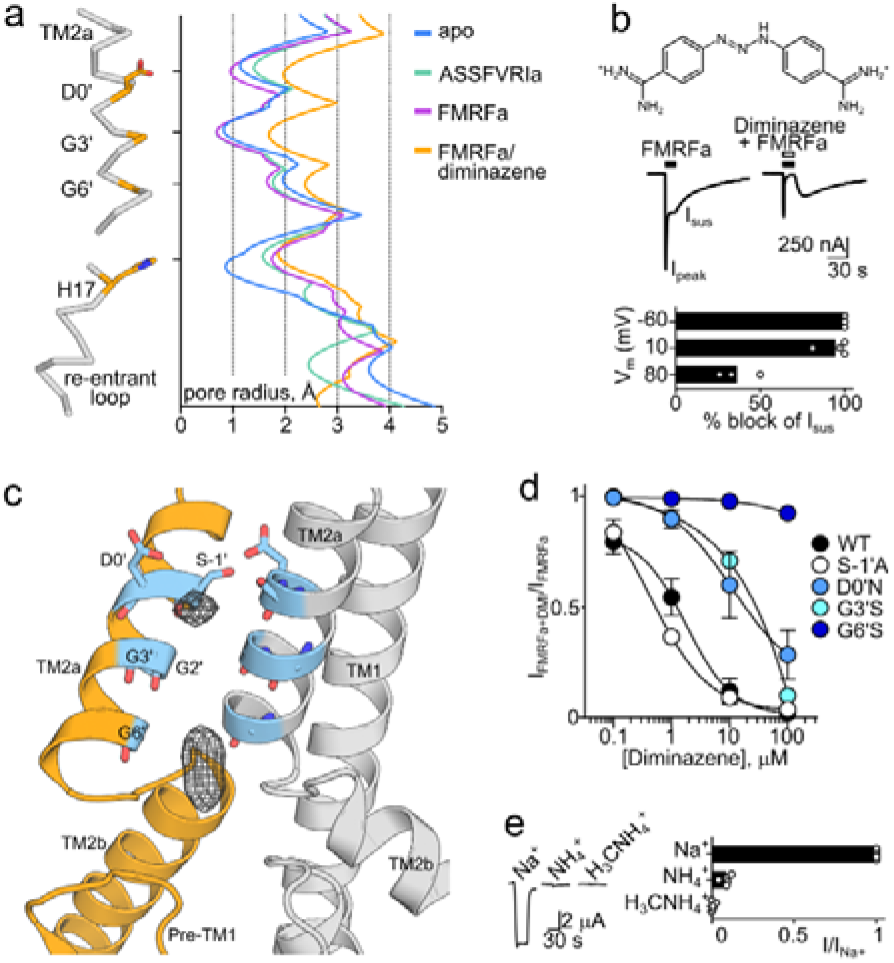
FaNaC1/FMRFa/diminazene structure. (a) Pore-lining residues in FaNaC1/FMRFa/diminazene structure (left) and pore radius for all four cryo-EM structures calculated in HOLE^73^ (right). (b) *Upper*, diminazene structure (ionized at pH 7.4). *Mid*, peak (I_peak_) and sustained (I_sus_) FMRFa (3 μM) -gated currents through FaNaC1 in the absence and presence of diminazene (10 μM). *Lower*, % block of I_sus_ by diminazene at three different membrane potentials. (c) Magnified view of FaNaC1/FMRFa/diminazene model, showing non-protein cryo-EM density in the pore as mesh. (d) Concentration-dependent block of I_sus_ by diminazene at wildtype (WT) and mutant channels (mean ± SEM, n = 4). (e) FMRFa-gated current amplitude (V_m_ = -60 mV) in extracellular solutions based on different cations.

We therefore sought structural data on a ligand-bound, open-channel state by solving the structure of FaNaC1 in the presence of both FMRFa and diminazene, a pore blocker of diverse DEG/ENaC channels^20,29,30^ that delays desensitization in ASICs by plugging the open channel pore^30^. We first verified that diminazene blocks FaNaC1 expressed in Xenopus oocytes by co-applying FMRFa and diminazene and observed inhibition of both peak (IC_50_ = 9 ± 3 μM, n = 5) and sustained FMRFa-gated current (IC_50_ = 1.8 ± 0.9 μM, n = 6; Fig. 4b, Fig. S6). Diminazene block was stronger at negative membrane potentials, and a large rebound current was observed after the removal of FMRFa and diminazene (Fig. 4b), suggesting that the positively charged drug inhibits FaNaC1 by plugging the open-channel pore and preventing the channel closure that occurs in desensitization.

In solving the FaNaC1/FMRFa/diminazene structure, several 3D classes emerged during our image analysis, with two predominating: a closed-channel class similar to the FaNaC1/FMRFa structure; and a class that differed from the others with dilated mid- to upper-pore, and some likely non-protein density in the channel pore (Fig. 4c, Fig. S7). We focused on this second 3D class, resolving the structure to 3 Å resolution (Fig. S7). We easily modelled TM2 helices into the cryo-EM density and observed a ∼1 Å increase in the pore diameter relative to our other structures (Fig. 4a). Although the resolved density in the pore was too small to accommodate a full diminazine molecule (Fig. 4b,c), no such density or dilated conformation was observed in the diminazine-free FMRFa-bound dataset (Fig. S2), suggesting that diminazene binds and affects FaNaC1 pore conformation.

Given the drug’s effect on pore diameter and its voltage-dependent block of currents, we hypothesized that this non-protein density derives from partially unresolved or low occupancy diminazene molecules. To investigate this further, we measured diminazene block of mutant FaNaC1 channels carrying single amino acid substitutions around this site. Increasing side chain volume around the density via the G6’S substitution drastically reduced diminazene potency, and a similar G3’S substitution one helical turn higher caused a more modest reduction in potency (Fig. 4c,d, Fig. S6), suggesting that diminazene binds intimately at the level of G6’. We also generated mutant G2’S channels but saw no currents in oocytes injected with these RNAs (n = 8 over two batches of ooyctes). An additional helical turn higher, the S-1’A substitution caused constitutive current, as expected for mutating the TM2a -1’ degenerin position^31^, but had no effect on diminazene potency (Fig. 4e, Fig. S6). In contrast, the D0’N substitution decreased diminazene potency ∼10-fold (Fig. 4e), suggesting that the relatively well conserved TM2a 0’ carboxylate is important for sensitivity to diminazene. This may explain why ENaCs, which instead possess asparagine at the 0’ position, are insensitive to diminazene^32^ and closely reflects computational docking of the drug to ASIC1, where the D0’ carboxylate engages the upper positively charged amidine moiety^30^. Thus, we interpret our FaNaC1/FMRFa/diminazene structure as an FMRFa-gated, diminazene-blocked, open-channel conformation.

Based on the pore radius of ∼2 Å at G3’, G6’ and the re-entrant loop in our putatively open-channel structure (Fig. 4a), FaNaC1 presumably passes partly dehydrated Na^+^ ions and its pore is narrower than the previously captured open-channel DEG/ENaC structure of ASIC1^11^, although the re-entrant loop was not resolved in the latter. Furthermore, we found that larger nitrogen-based cations, ammonium and methylammonium, were much less permeant than Na^+^ in FaNaC1 (Fig. 4f), as observed for ENaC^33^ but different from ASIC1, which conducts substantial ammonium and methtylammonium current^34,35^. This suggests that FaNaC1 adopts a narrower open-channel pore than ASIC1, as represented by our FMRFa-gated, diminazene-blocked structure, and that FaNaC1 presumably passes partly dehydrated Na^+^ ions.

### Ligand-induced channel gating

Finally, we compared ligand-free FaNaC1 and FaNaC1/FMRFa/diminazene structures to establish the structural mechanism by which FMRFa binding opens the channel. In its binding pocket, FMRFa draws the C-terminal end of α2 and α3a (distal finger domain) three to five Å upward and inward in the direction of α6 of the adjacent subunit, whereas the N-terminal end of α2, α3b/α3c, and all of α6 (knuckle), are relatively static (Fig. 3d, Fig. 5a,b). This in turn draws the peripheral and long, vertical α5 segment (thumb) ∼3 Å inwards and ∼2 Å upwards (Fig. 5a,b). In contrast to the peripheral thumb, the more internal loops and β-sheets of the palm and ball domains remain relatively static, exemplified by F74 and Y174 side chains in a hydrophobic hub within this region (Fig. 5b). The net result of peripheral movement on internal stasis is that the extracellular domain of a single subunit rolls anti-clockwise (viewed from above) and upwards (viewed from the side, Fig. 5c). Extracellular domain rolling pulls the β-turn at the proximal-thumb/wrist domain outwards, which is coupled to the upper-channel expansion via β-turn H297 interactions with TM1 Y59 and TM2a E490 (Fig. 5b,c).

**Figure 5.**
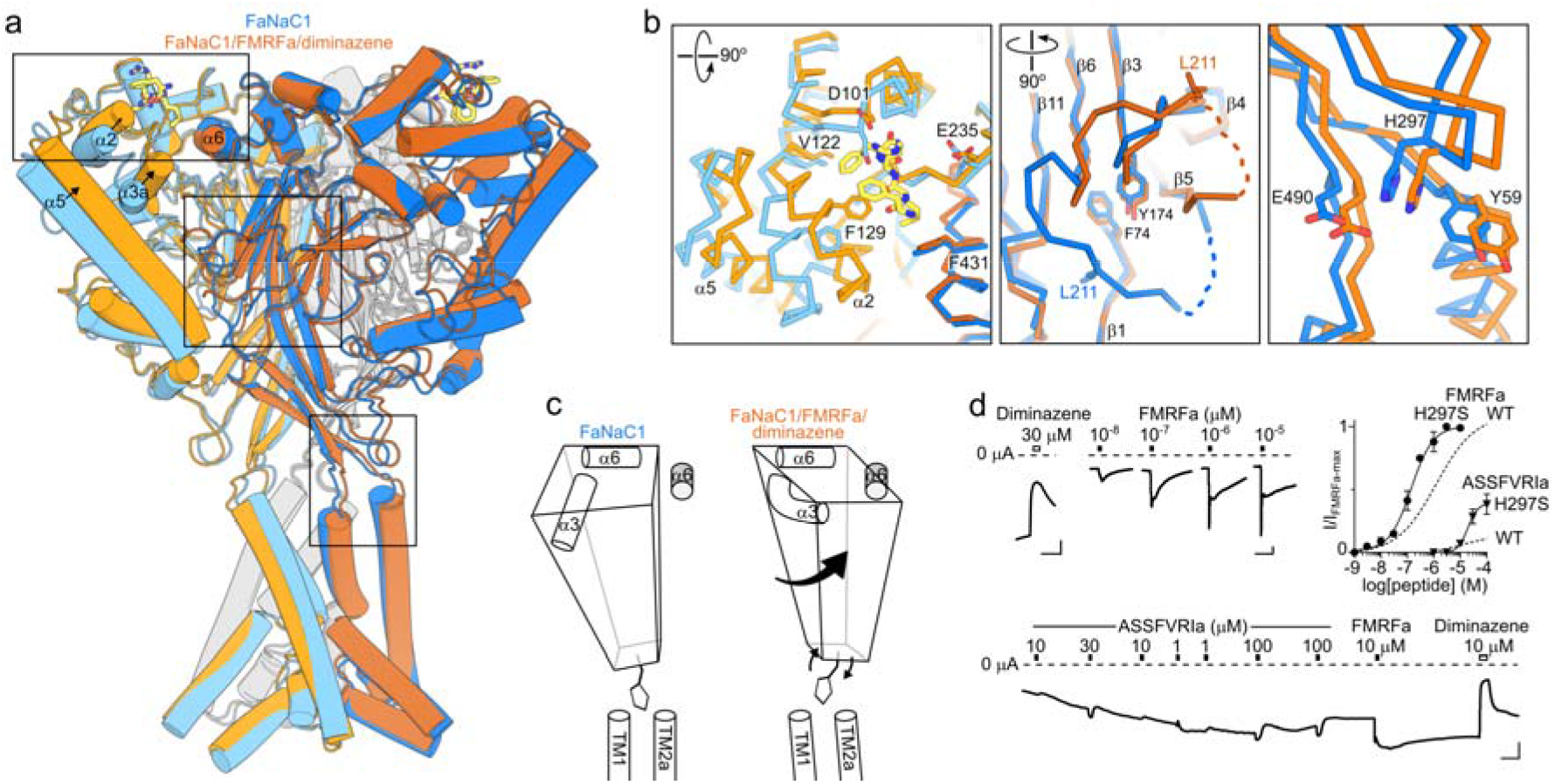
FaNaC1 gating mechanism. (a) Overlay of ligand-free FaNaC1 (blue) and FaNaC1/FMRFa/diminazene (orange) structures. Adjacent subunit in darker shades. (b) Magnified views of boxed regions, left to right: ligand binding site; palm domain; β-turn—TM2a/TM1 interactions. Selected side chains shown as sticks and labelled. (c) Illustration of one subunit (left subunit in **a**) moving during gating: peripheral rolling, internal stasis, and β-turn moving outward to allow channel expansion. α6 helix from adjacent subunit in grey. (d) *Upper-left and lower*, example recordings from oocytes expressing mutant H297S FaNaC1 channels (scale bars: x, 30 s; y, 1 μA). *Upper-right*, mean (± SEM, n = 5) responses of H297S channels to FMRFa and ASSFVRIa normalized to maximum FMRFa-gated current (I/I_max-FMRFa_), compared to WT channels.

Seeking verification that β-turn—TM1/TM2 interactions mediate channel gating, we tested the activity of mutant H297S channels in which the large H297 side chain is replaced with a much smaller side chain, making these interactions less likely. Remarkably, H297S channels were constitutively active, with 3.8 ± 1.3 μA (n = 10) current in the absence of agonist (cf. 120 ± 40 nA in WT, n = 10), which was blocked on average 76 ± 5% (n = 7) by 30 μM diminazene (Fig. 5d). Furthermore, the addition of FMRFa activated additional current, with increased potency compared to WT, and the relative efficacy of partial agonist ASSFVRIa was increased by the mutation (Fig. 5d). Thus, the H297S mutation increases gating efficacy. This suggests that wildtype resting FaNaC1 channels are energetically primed for opening but cannot do so until ligand-binding and extracellular domain-rolling releases TM1/TM2a via outward H297 movement.

We find that ligand-induced channel gating extends to the lower part of the pore, observing a 1 Å increase in pore radius at the level of re-entrant loop T15 and H17 side chains compared to the ligand-free state (Fig. 4a). This offers structural evidence for lower-pore gating that was suggested to occur in molluscan FaNaC and mammalian ENaC based on electrophysiological studies^36-38^. Lower-pore dilation is similar in both FaNaC1/FMRFa/diminazene and FaNaC1/FMRFa structures (Fig. 4a), in contrast to upper-pore dilation, which is present in the FaNaC1/FMRFa/diminazene structure but collapsed in the desensitized FaNaC1/FMRFa structure (Fig. 4a). This suggests that desensitization is an upper-channel and extracellular vestibule phenomenon. We also observed a remarkably large conformational change in the β5-β6 loop (residues K200-G218) of the extracellular domain, which flips from a downward orientation in our ligand-free structure to an upward orientation in all ligand-bound structures (Fig. 5a,b). This involves, for example, L211 displacing 20 Å (Fig 5b) and is a notable exception to the otherwise immobile palm/ball domain. Whether β5-β6 loop flipping is a consequence of activation or a diminazine-insensitive aspect of desensitization would require further investigation.

## DISCUSSION

### Divergent ligand-recognition in the DEG/ENaC superfamily

Our four Malacoceros FaNaC1 structures offer a precise description of neuropeptide binding and a comprehensive view of ligand-induced channel gating in a DEG/ENaC channel. The extracellular neuropeptide-binding pocket is formed by dynamic α1-α3 segments of the finger domain and more static β6-β7 and α6 segments of proximal-finger and knuckle domains. Comparison with other DEG/ENaCs show similar overall architecture but divergent sequence and secondary and tertiary structure in the first three helices of the finger domain (Fig. S7). Despite its divergence in different DEG/ENaCs, this external corner of the DEG/ENaC trimer appears to play important roles in gating throughout the superfamily. It includes modulatory ion- and protease-binding sites in ENaC^15,42^, residues whose mutation decreases pH sensitivity in ASICs^43^, and proposed sites for extracellular matrix tethering to mechano-sensitive DEG/ENaCs^44^, in addition to residues in mollusc FaNaCs whose mutation decreases FMRFa potency^24,45,46^.

We were surprised to find, however, that even within the FaNaC family, residues of the Malacoceros FaNaC1 FMRFa-binding pocket are poorly conserved. Although the α2-122 residue (V122 in Fig. 2a,d) is relatively well conserved (valine in FaNaC1; valine or isoleucine in most FaNaCs), α1, α2, and α3a sequence is divergent and difficult to align (Fig. S8). This includes α2-F129, the most influential residue for FMRFa potency in our experiments, which seems to be restricted to annelid FaNaCs, although mollusc FaNaCs also possess putative-α2 phenylalanine residues (Fig. 2d), and a potentially homologous role in ligand recognition is supported by the decrease in FMRFa caused by their substitution in a recent study^24^. Nonetheless, some divergence is to be expected, as annelid FaNaCs are gated by partial agonists ASSFVRIa and LFRYa^6^, in contrast to mollusc FaNaCs, which are instead gated by partial agonists more closely analogous to FRMFa^6,19,25,26^. The more strictly conserved determinants of agonist potency we previously identified throughout the FaNaC family are in a different site, between palm domains of adjacent subunits {Dandamudi, 2022 #454}. These are likely important for coupling divergent ligand binding in the finger domain to channel gating below.

### Channel gating and ion conduction illuminated by FaNaC1 structures

Previously, our understanding of DEG/ENaC gating mechanisms and ion conduction was based on chicken ASIC1 structures in resting, active, and desensitized states^11,12,23^. The FaNaC1 structures presented here capture a ligand-free resting state, two ligand-bound desensitized states, and a ligand-bound dilated channel state. The comparison of these structures points to a gating mechanism in which the outer finger domain closes around the agonist, starting an anticlockwise rotation of the extracellular domain (Fig. 5). As the palm remains static, the periphery of the extracellular domain (the thumb) rolls anti-clockwise and upwards, pulling the upper part of the channel (wrist) outwards, resulting in pore dilation. This anticlockwise extracellular domain rotation and wrist expansion loosely reflects gating of ASIC1^12^, although the principal trigger(s) initiating these conformational changes in ASIC1 are yet to be identified^13^. One major difference between FaNaC1 and ASIC1 gating regards desensitization. The large rearrangement of the β11-β12 linker during ASIC1 desensitization^12^ is not observed in FaNaC1, although we do observe a large conformational change of the FaNaC1 β5-β6 loop, which is close to the β11-β12 linker (Fig. 5b).

A major advance of our study is the capture of a potentially open-channel pore structure. In previous high-resolution DEG/ENaC structures the channel domain was difficult to resolve^11,12,15^, and in particular, the only open-channel structure of chicken ASIC1 does not include the pore-lining pre-TM1 re-entrant loop^11^. Nonetheless, a better resolved picture of ion conduction throughout the superfamily emerges from the comparison of FaNaC1 and distantly related ASIC1 pores. FaNaC1 is more closely related to ENaC^8,47,48^, both of these channels are poorly permeable to nitrogen-based cations larger than Na^+ 33^, and our FaNaC1/FMRFa/diminazene structure reveals an ion pathway slightly narrower than both ASIC1 and voltage-gated sodium channels^11,49^, both of which pass ammonium, methylammonium, and hydroxylamine relatively well^34,35,50^. Taken together, our capture of FaNaC1 in different functional states describes the mechanism by which FMRFa elicits excitatory neuronal signals and offers a template for future studies dissecting channel gating and ion conduction in DEG/ENaCs.

## METHODS

### Cell lines

Adherent HEK293T cells were cultured in 10 cm Petri dishes in DMEM with L-glutamine and sodium pyruvate (Gibco), supplemented with 10% FBS and antibiotic-antimycotic at 37 °C and 5% CO_2_. Suspension HEK293S GnTI^-^ cells were maintained in Freestyle media with GlutaMAX (Gibco) supplemented with 1% FBS and antibiotic-antimycotic solution, at 37 °C, 5% CO_2_ and 60% humidity, in TPP600 bioreactors. Sf9 cells were cultured in SFMIII media supplemented with antibiotic-antimycotic, at 27 °C.

### Protein expression and purification

Commercially synthesized Malacoceros fuliginosus FaNaC1 and Octopus bimaculoides FaNaC sequences^6^ (modified from Genbank entries ON156825 and XM_014930938.1 to remove internal restriction sites) were cloned into a pEZT-BM vector^51^ adapted for FX-cloning^52^ with C-terminal HRV-3C cleavage site, Venus YFP, myc and SBP-tags. For homolog screening, adherent HEK293T cells at ∼60% confluency were transfected with these plasmids using PEI 40K MAX (DNA:PEI ratio of 1:3, 10 ug DNA/dish). Proteins were expressed for 48 h, cells were harvested, washed with PBS and stored at -80 °C until further use. Cells from 1 dish were resuspended in 200 ul extraction buffer (2% DDM, 0.4% CHS, 20 mM HEPES pH 7.6, 150 mM NaCl, 10% glycerol, cOmplete protease inhibitors), proteins were extracted for 2 h. Lysate was centrifuged at 150 000 g and supernatant analyzed on Tosoh G4000PWXL using FSEC^53^.

Large-scale expression of Malacoceros FaNaC1 was performed using BacMam expression system^54^. Viruses were generated as described^51,54^. Briefly, the Malacoceros FaNaC1 bacmid was generated following the Invitrogen Bac-to-Bac protocol. Afterwards, Sf9 cells were transfected with the bacmid using Cellfectin according to manufacturer’s instructions. 4-5 days after transfection, the P0-containing supernatant was harvested, and supplemented with 10% FBS, and this virus stock was used to generate P1. Sf9 cells were infected at density 1 × 10^6^, and once the majority of cells were fluorescent, the supernatant containing virus was filtered, supplemented with 10% FBS and stored at 4 °C until further use. One day prior to infection, HEK293S were split to 0.6 × 10^6^ cells/ml density. The following day, titerless P1 virus was diluted 1:10 into expression culture. After 24 h, sodium butyrate was added to a final concentration of 10 mM and the protein was expressed for additional 48 h. Cells were harvested, washed with PBS and stored at -80 °C.

All of the purification steps were performed on ice or at 4 °C. Cell pellets from ∼6 L of expression culture were resuspended in buffer A (20 mM HEPES pH 7.6, 150 mM NaCl, 10% glycerol, DNAse I, 2 mM MgCl2, 2% DDM, 0.4% CHS, cOmplete protease inhibitor tablets), and the protein was extracted for 2 h under gentle agitation. The lysate was centrifugated at 200,000 g for 30 min, the supernatant was incubated with the affinity resin with immobilized anti-GFP nanobody 3K1K^55^ for 30 min, and passed through the resin 3-4 times in a gravity flow column. The resin was washed with ∼30 column volumes (CV) of buffer B (20 mM HEPES pH 7.6, 150 mM NaCl, 10% glycerol, 0.02% GDN). Afterwards, the protein was cleaved off the resin in batch with HRV-3C protease (∼1.2 mg) for 2 h. The eluate was concentrated using 100 kDa cut-off Amicon centrifugal filter units at 600 g and injected onto a Superose 6 Increase 10/300 column equilibrated in buffer C (20 mM HEPES pH 7.6, 150 mM NaCl, 0.02% GDN). Main peak fractions were pooled and concentrated as described above.

### Nanodisc reconstitution

Nanodisc reconstitution was performed as described^56^. Briefly, lipids (POPC:POPG 3:1 molar ratio) were pooled, dried using rotary evaporator, and washed with diethyl ether. After diethyl ether was evaporated, the lipids were rehydrated in ND buffer (20 mM HEPES, 150 mM NaCl, 30 mM DDM) at a concentration of 10 mM. Purified protein was mixed with lipids, incubated for 30 minutes, after which the purified MSP was added, incubated for 30 min, followed by addition of SM-2 biobeads (200 mg/ml of assembly reaction). The assembly ratios FaNaC:lipids:MSP were 3:1100:10, assuming 1 FaNaC trimer per 5 assembled nanodiscs. The mixture was incubated overnight at 4 degrees with gentle agitation. The following day, the sample was concentrated in 100 kDa Amicon concentrators at 500 g and injected onto a Superose 6 Increase 10/300 column equilibrated in buffer D (20 mM HEPES pH 7.6, 150 mM NaCl). Higher MW peak containing nanodisc-reconstituted FaNaC1 was pooled and concentrated as above to 1.4 – 1.8 mg/ml. Ligands and diminazene were added directly before freezing (FMRFa 30 μM, ASSFVRIa 100 μM, diminazene 100 μM).

### Cryo-EM sample preparation and data acquisition

Quantifoil 1.2/1.3 Au grids with 300 mesh were glow-discharged at 5 mA for 30 s. The grids were frozen using Vitrobot Mark IV (Thermo Fisher). 2.8 ul of freshly prepared sample were applied to the grids, which were blotted for 3.5 s with blot force 0 at 15 °C and 100% humidity. The grids were plunge-frozen in ethane-propane mixture and stored in liquid nitrogen until further use. The data were recorded at the University of Groningen on a 200 keV Talos Arctica (Thermo Fisher) with a K2 summit detector (Gatan), post-column energy filter, 20 eV slit and a 100 μm objective aperture. Optimal squares for data collection were selected using an in-house sample thickness estimation script^57^. The images were recorded in an automated fashion using SerialEM^58^ with a 3×3 multishot pattern. Cryo-EM images were acquired at a pixel size of 1.022 Å (calibrated magnification of 49 407x), a defocus range from –0.5 to –2 μm, an exposure time of 9 s with a sub-frame exposure time of 150 ms (60 frames), and a total electron exposure on the specimen of about 52 electrons per Å^2^. Micrographs were pre-processed on the fly in FOCUS^59^ using MotionCor2^60^ for motion correction and ctffind4.1^61^ for contrast transfer function (CTF) resolution estimation. Images with the defocus values 0.4 – 2 μm, showing no ice contamination and with a CTF resolution estimate better than 6 Å were selected for further processing.

### Image processing

The collected datasets were processed following an essentially identical scheme, with the exception of the FaNaC1/FMRFa/diminazene dataset. Particles were picked using a general model in crYOLO^62^, and subsequently extracted in Relion-3.1^63^ with a box size of 220 pixels for FMRFa, ASSFVRIa, and FMRFa/diminazene datasets, and 240 for apo, respectively. The extracted particles were imported into cryoSPARC^64^ and subjected to 2D classification (initial batch size 200, 10 final full iterations). Particles from selected 2D classes were subjected to ab initio 3D reconstruction with 5 classes and subsequent heterogeneous refinement using all 5 ab initio classes as an input. Particles from the best class were imported into Relion-3.1, and subjected to Bayesian polishing followed by several rounds of CTF refinement. In the case of the diminazene-supplemented dataset, particles were further classified with no image alignment and a mask covering the transmembrane part in order to resolve the conformation heterogeneity in the pore region. In all cases, the final sets of particles were subjected to masked refinement with a C3 symmetry imposed. The half-maps were used as inputs for postprocessing in deepEMhancer^65^ with a tight model.

### Model building and refinement and pore analysis

The initial model of Malacoceros FaNaC1 was predicted using Alphafold^66^ and adjusted manually in coot^67^. The ligand coordinates were generated in Chem3D 18 (PerkinElmer), and the restraints for the ligands were generated in PRODRG^68^. Models were iteratively adjusted in coot and ISOLDE^69^, followed by real-space refinement in Phenix^70^ with NCS and secondary structure restraints against a refined unsharpened map. Figures were prepared in Pymol, Chimera^71^ and ChimeraX^72^. Channel pore radius was calculated with HOLE^73^.

### Electrophysiology and data analysis

FaNaC1 in a modified pSP64 plasmid vector was used for mutagenesis and mRNA preparation and injection into stage V/VI Xenopus laevis oocytes (EcoCyte Bioscience, Germany) as described previously vector^6^. Mutants were generated by partly overlapping primers^74^, and channel-coding inserts were Sanger sequenced. FMRFamide and ASSFVRIamide (acetate salts, custom synthesized by Genscript, purity 95.1-99.6% purity by HPLC, mass confirmed by ESI-MS) were dissolved in water to 10 mM and diminazene aceturate (Merck) was dissolved in DMSO to 100 mM before dilution in experimental solution: (in mM) NaCl 96, KCl 2, CaCl_2_ 1.8, MgCl_2_ 1, HEPES 5, pH 7.5 (NaOH). NaCl was replaced with KCl, NH_4_Cl, or H_3_CNH_3_Cl (Merck) where appropriate. Oocytes were clamped at -60 mV unless otherwise indicated.

Currents were measured by two electrode voltage clamp, as previously described^6^, using a Warner OC-725C amplifier and HEKA LIH8+8 interface, sampling at 500 Hz or 1 kHz and filtering at 100 Hz. Current amplitudes were measured in Clampfit 11 (Molecular Devices). Data were analyzed in Prism 9 (Graphpad) and fit with Prism 9 variable-slope 4-parameter nonlinear regression yielding half-maximal effective activating/inhibiting concentrations (EC_50_/IC_50_). Mean ± SEM EC_50_/IC_50_ from fits to individual cells reported in main text, fits to averaged data points shown in figures. Currents shown in figures are further filtered (20 Hz) and decimated (50 x) in Clampfit 11 for smaller filesize.

## ACKNOWLEDGEMENTS

VK was supported by SNSF Postdoc.Mobility Fellowship P500PB_203053, CP by the Dutch Research Council (NWO) grant 722.017.001 and 740.018.016, TL by The Research Council of Norway, project number 234817. The authors would like to thank J. Rheinberger and M. Punter for facility and image processing cluster management. Members of Paulino and Lynagh labs are acknowledged for technical help and discussions at various stages of the project.

## AUTHOR INFORMATION

VK, project conception and design, molecular biology, protein expression and purification, cryo-EM data collection and analysis, figure preparation, manuscript editing. MD, electrophysiology experiments and analysis, figure preparation, manuscript editing. CP, supervision, manuscript editing. TL, project design, supervision, molecular biology, electrophysiology experiments and analysis, figure preparation, manuscript writing.

