## Supplementary material for "Structural basis for excitatory neuropeptide signaling": Table S1 and Figures S1-S8

[#] Current address, Department of Biomedicine, University of Bergen, Bergen, Norway

**This file contains one table and seven figures.**

Table S1. Cryo-EM data collection, refinement and validation statistics

Figure S1. Ligand-free FaNaC1 structure determination by cryo-EM

Figure S2. FMRFa-bound FaNaC1 structure determination by cryo-EM

Figure S3. Effect of FaNaC1 on the surrounding lipid bilayer

Figure S4. Comparison of FaNaC1 and other DEG/ENaC structures.

Figure S5. ASSFVRIa-bound FaNaC1 structure determination by cryo-EM

Figure S6. Effects of diminazene on mutant FaNaC1 channels

Figure S7. FMRFa-bound FaNaC1 in the presence of diminazene, structure determination by cryo-EM

Figure S8. Amino acid sequence alignment of FaNaCs and other DEG/ENaCs

**Table S1. Cryo-EM data collection, refinement and validation statistics**

| | Apo (EMDB-16982) (PDB 8ON8) | FMRFa (EMDB-16981) (PDB 8ON7) | ASSFVRIa (EMDB-16983) (PDB 8ON9) | FMRFa / dim. (EMDB-16984) (PDB 8ONA) |
|---|---|---|---|---|
| **Data collection and processing** | | | | |
| Magnification | 49,407 | 49,407 | 49,407 | 49,407 |
| Voltage (kV) | 200 | 200 | 200 | 200 |
| Electron exposure (e–/Å$^2$) | 47.47 | 50.11 | 47.40 | 47.20 |
| Defocus range (μm) | -0.3 to -2.0 | -0.3 to -2.0 | -0.3 to -2.0 | -0.3 to -2.0 |
| Pixel size (Å) | 1.022 | 1.022 | 1.022 | 1.022 |
| Symmetry imposed | C3 | C3 | C3 | C3 |
| Initial particle images (no.) | 2.659.492 | 3.235.905 | 4.662.322 | 3.893.627 |
| Final particle images (no.) | 376.869 | 458.934 | 477.006 | 137.558 |
| Map resolution (Å) | | | | |
| FSC threshold 0.143 | 2.67 | 2.52 | 2.39 | 2.96 |
| Map resolution range (Å) | 2.6-3.5 | 2.4-3.6 | 2.3-3.5 | 2.9-3.8 |
| **Refinement** | | | | |
| Initial model used (PDB code) | 8ON7 | Alphafold | 8ON7 | 8ON7 |
| Model resolution (Å) | | | | |
| FSC threshold 0.143 | 2.4 | 2.2 | 2.2 | 2.7 |
| Model resolution range (Å) | 2.4-3.5 | 2.2-3.6 | 2.2-3.5 | 2.7-3.8 |
| Map sharpening *B* factor (Å$^2$) | N/A | N/A | N/A | N/A |
| Q-score | 0.49 | 0.55 | 0.53 | 0.50 |
| Model composition | | | | |
| Non-hydrogen atoms | 12891 | 13032 | 13011 | 12900 |
| Protein residues | 1572 | 1575 | 1572 | 1569 |
| Ligands | 21 | 24 | 24 | 18 |
| *B* factors (Å$^2$) | | | | |
| Protein | 118.56 | 96.61 | 104.67 | 123.87 |
| Ligand | 143.93 | 112.75 | 129.08 | 134.85 |
| R.m.s. deviations | | | | |
| Bond lengths (Å) | 0.002 | 0.003 | 0.006 | 0.004 |
| Bond angles (°) | 0.498 | 0.721 | 0.740 | 0.706 |
| Validation | | | | |
| MolProbity score | 1.19 | 1.33 | 1.41 | 1.32 |
| Clashscore | 3.76 | 5.03 | 7.51 | 5.83 |
| Poor rotamers (%) | 0.2 | 0.4 | 0.9 | 0.6 |
| Rama Z | | | | |
| Whole | 1.69 (0.22) | 0.45 (0.20) | 0.33 (0.20) | 0.68 (0.21) |
| Helix | 1.56 (0.20) | 0.17 (0.19) | 0.07 (0.18) | 0.20 (0.19) |
| Sheet | 1.75 (0.33) | 1.28 (0.29) | 1.42 (0.29) | 1.09 (0.30) |
| Loop | 0.45(0.27) | 0.22 (0.25) | 0.06 (0.25) | 0.71 (0.26) |
| Ramachandran plot | | | | |
| Favored (%) | 97.88 | 97.70 | 98.27 | 98.27 |
| Allowed (%) | 2.12 | 2.10 | 1.53 | 1.53 |
| Disallowed (%) | 0 | 0.2 | 0.2 | 0.2 |

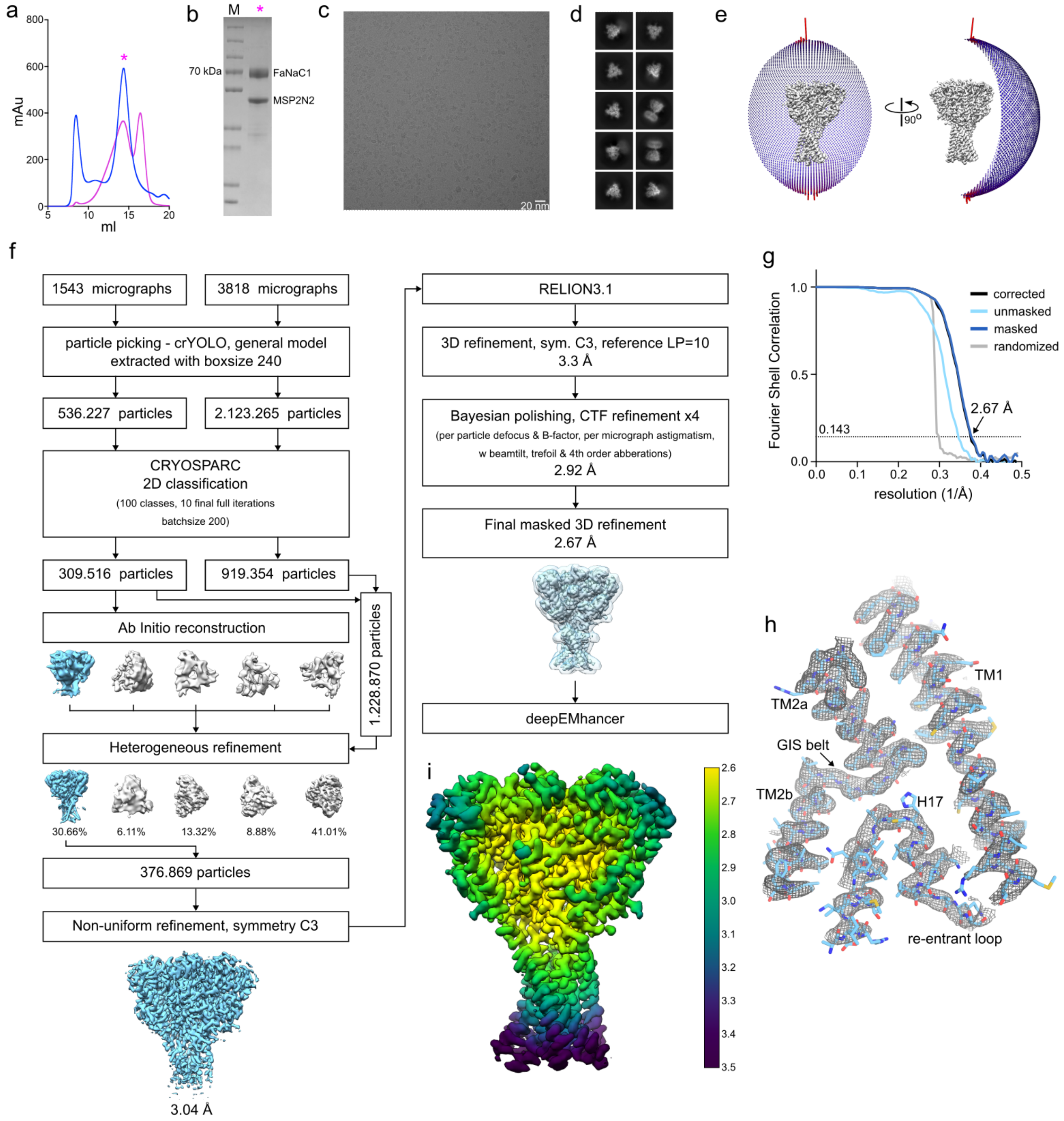


**Figure S1. Ligand-free FaNaC1 structure determination by cryo-EM**

(a) Size exclusion profiles of detergent-solubilised (blue) and nanodisc-reconstituted (magenta) *Malacoceros* FaNaC1. Samples were analysed on Superose 6 Increase 10/300 column. (b) SDS-PAGE of the main peak fraction, which was used for sample preparation for cryo-EM. Representative cryo-EM image (c) and 2D classes (d) of vitrified FaNaC1 in apo state. (e) Angular distribution of the particles included in the final C3-symmetrised map. The length and the colour of the sticks represent the number of particles. (f) Schematic representation of the processing workflow, the mask used in the final refinement iteration is displayed as a transparent outline. (g) FSC plot used for resolution estimation (0.143 cut-off criteria). (h) Density corresponding to the transmembrane domain shown as grey mesh, deepEMhancer map used for visualization is contoured at 4.5 σ. The respective fitted model is shown in blue. I. Final deepEMhancer-postprocessed map coloured according to the local resolution estimation in Relion.

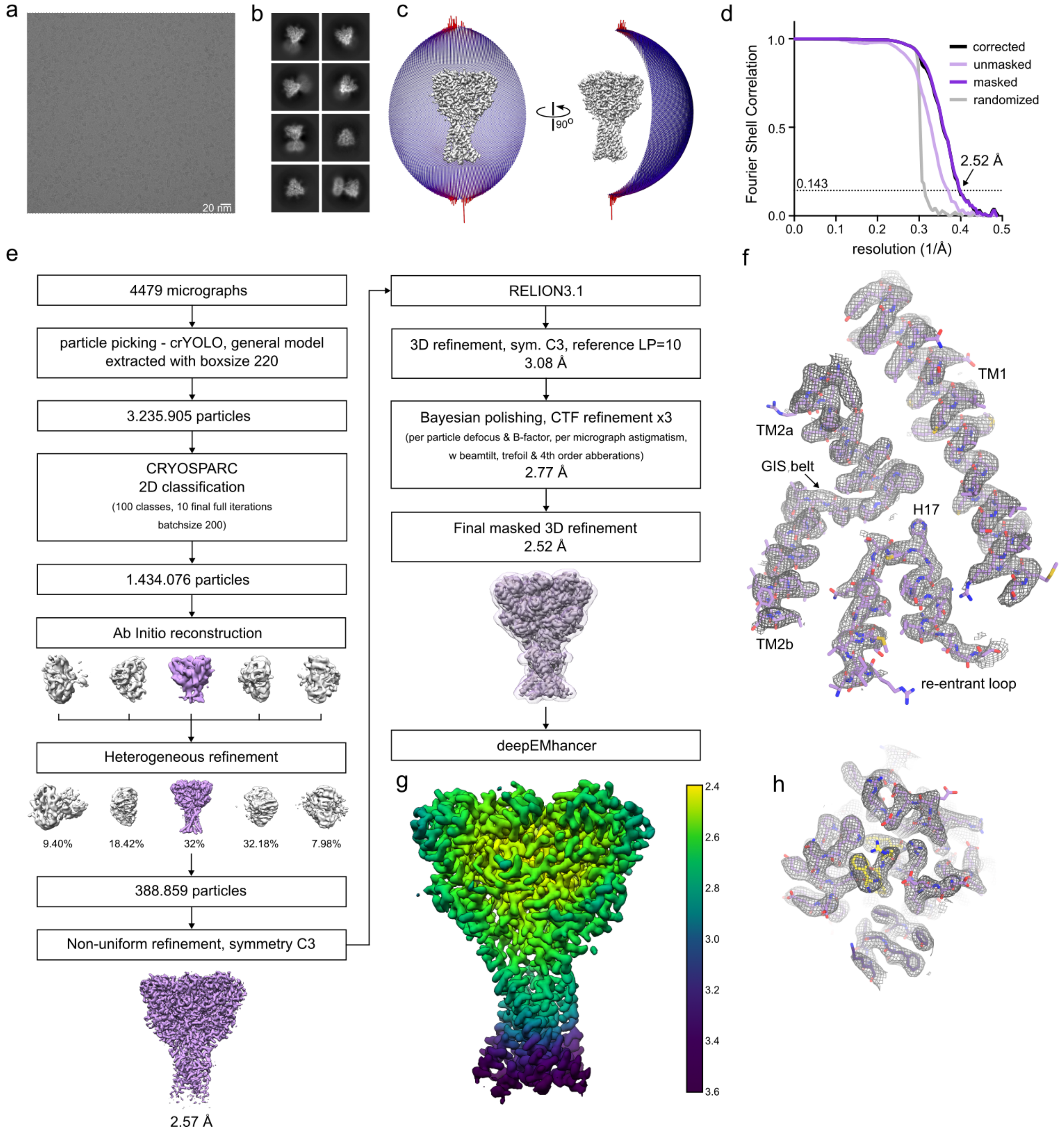


**Figure S2. FMRFa-bound FaNaC1 structure determination by cryo-EM**

Representative cryo-EM image (a) and 2D classes (b) of vitrified FaNaC1 in presence of FMRFa. (c) Angular distribution of the particles included in the final C3-symmetrised map. The length and the colour of the sticks represent the number of particles. (d) FSC plot used for resolution estimation (0.143 cut-off criteria). (e) Schematic representation of the processing workflow, the mask used in the final refinement iteration is displayed as a transparent outline. (f) Density corresponding to the transmembrane domain shown as grey mesh, deepEMhancer map used for visualization is contoured at 4.5 σ. The respective fitted model is shown in purple. (g) Final deepEMhancer-postprocessed map coloured according to the local resolution estimation in Relion. (h) Close-up view of the ligand-binding site, deepEMhancer map is displayed as grey mesh contoured at 4.5 σ, fitted atomic model is displayed in purple, FMRFa molecule is shown in yellow.

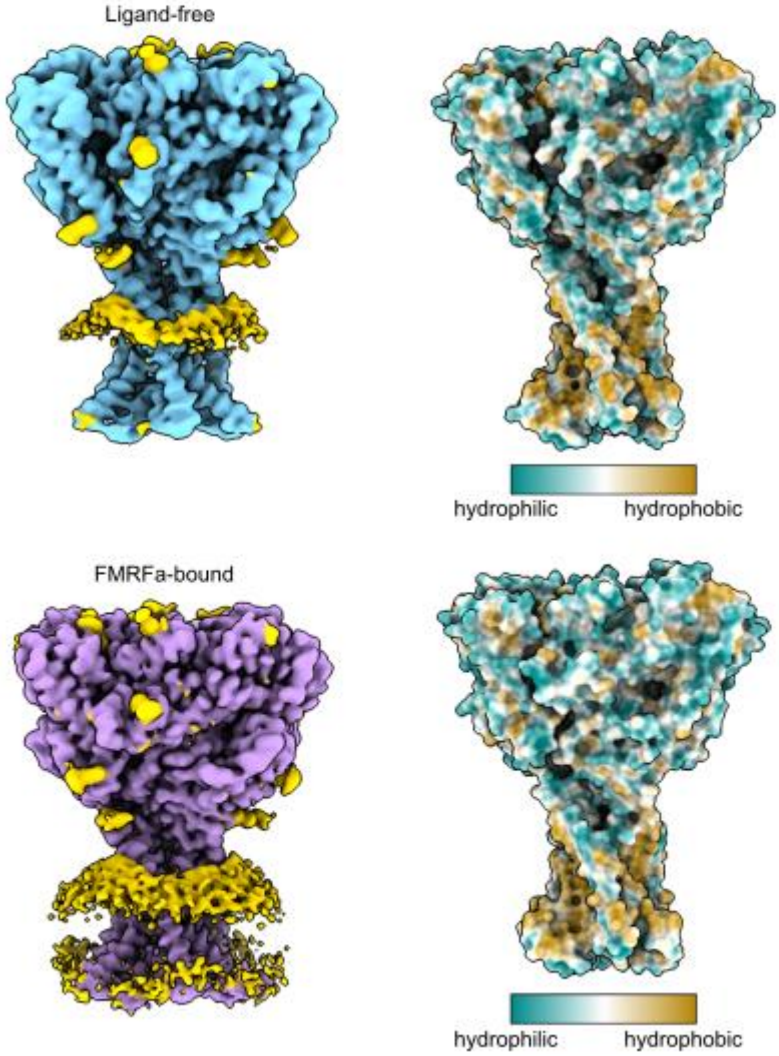


**Figure S3. Effect of FaNaC1 on the surrounding lipid bilayer.** Shown are unmasked refined maps of FaNaC1 in apo- (blue) and FMRFa-bound (purple) states, density corresponding to nanodisc and glycosylation sites are shown in yellow. Right, surface representation of the refined models coloured according to hydrophobicity.

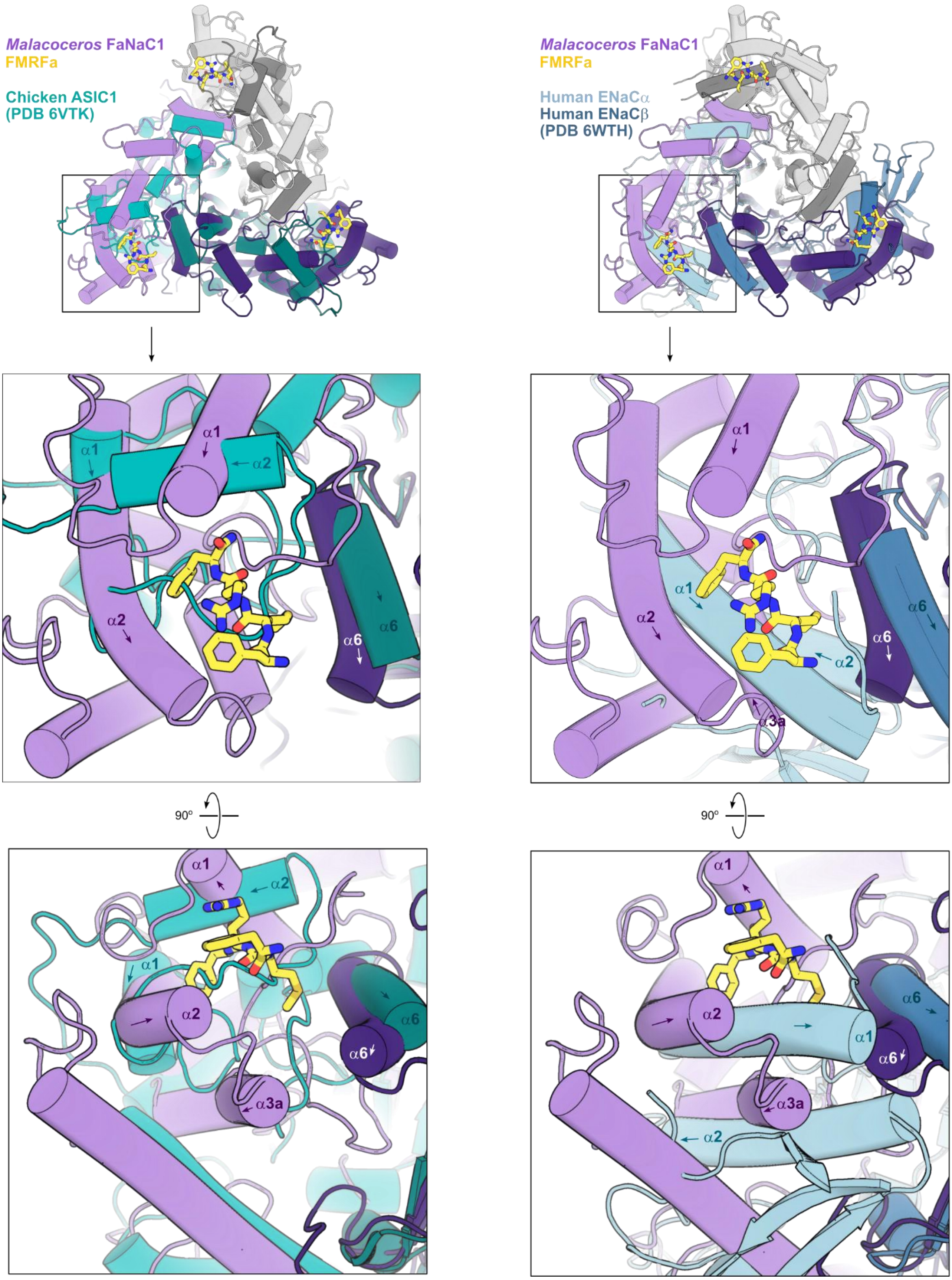


**Figure S4. Comparison of FaNaC1 and other DEG/ENaC structures.**

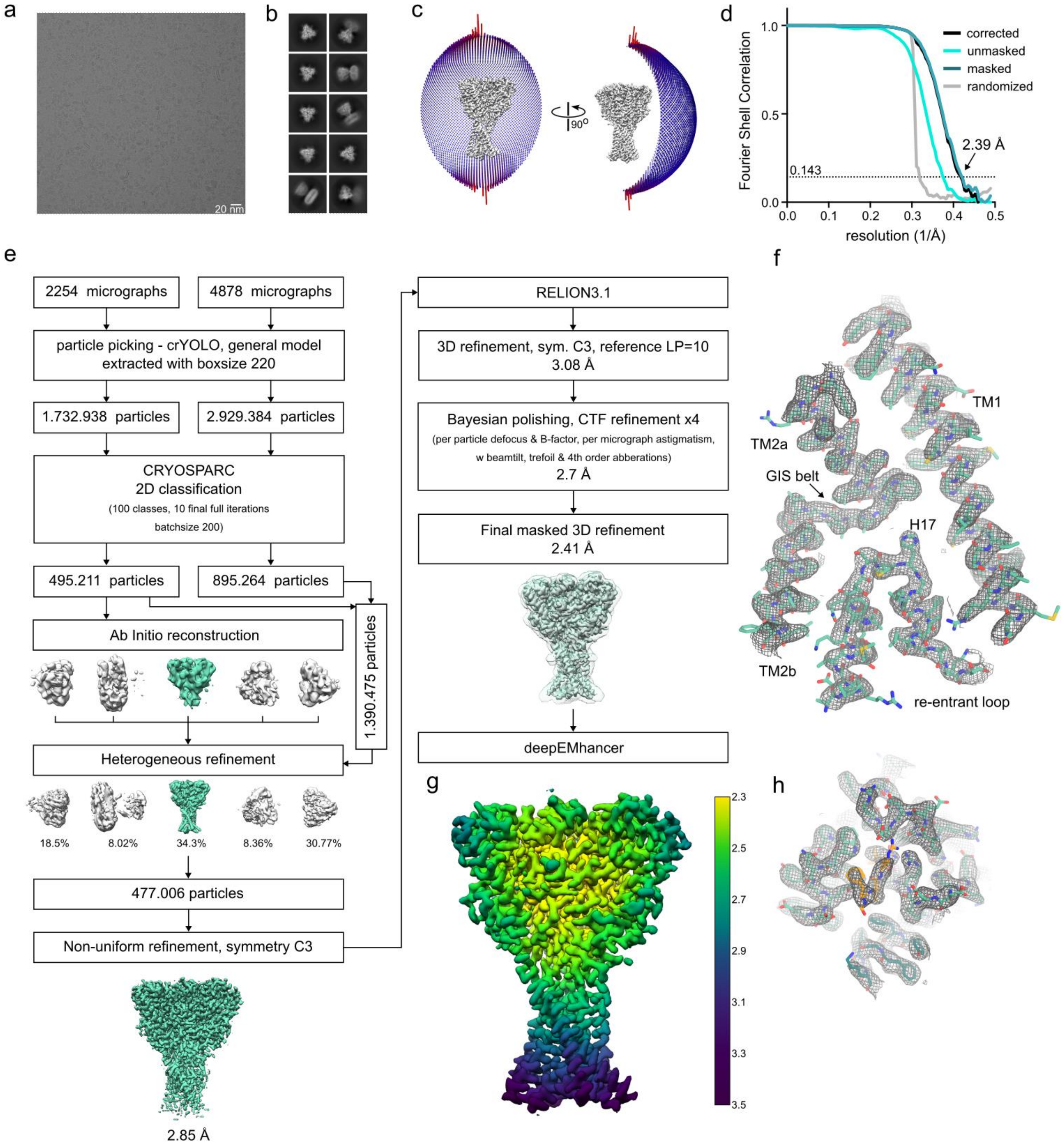


**Figure S5. ASSFVRIa-bound FaNaC1 structure determination by cryo-EM**

Representative cryo-EM image (a) and 2D classes (b) of vitrified FaNaC1 in ASSFVRIa-bound state. (c) Angular distribution of the particles included in the final C3-symmetrised map. The length and the colour of the sticks represent the number of particles. (d) FSC plot used for resolution estimation (0.143 cut-off criteria). (e) Schematic representation of the processing workflow, the mask used in the final refinement iteration is displayed as a transparent outline. (f) Density corresponding to the transmembrane domain shown as grey mesh, deepEMhancer map used for visualization is contoured at 4.5 σ. The respective fitted model is shown in teal. (g) Final deepEMhancer-postprocessed map coloured according to the local resolution estimation in Relion. (h) Close-up view of the ligand-binding site, deepEMhancer map is displayed as grey mesh contoured at 4.5 σ, fitted atomic model is displayed in teal, FMRFa molecule is shown in orange.

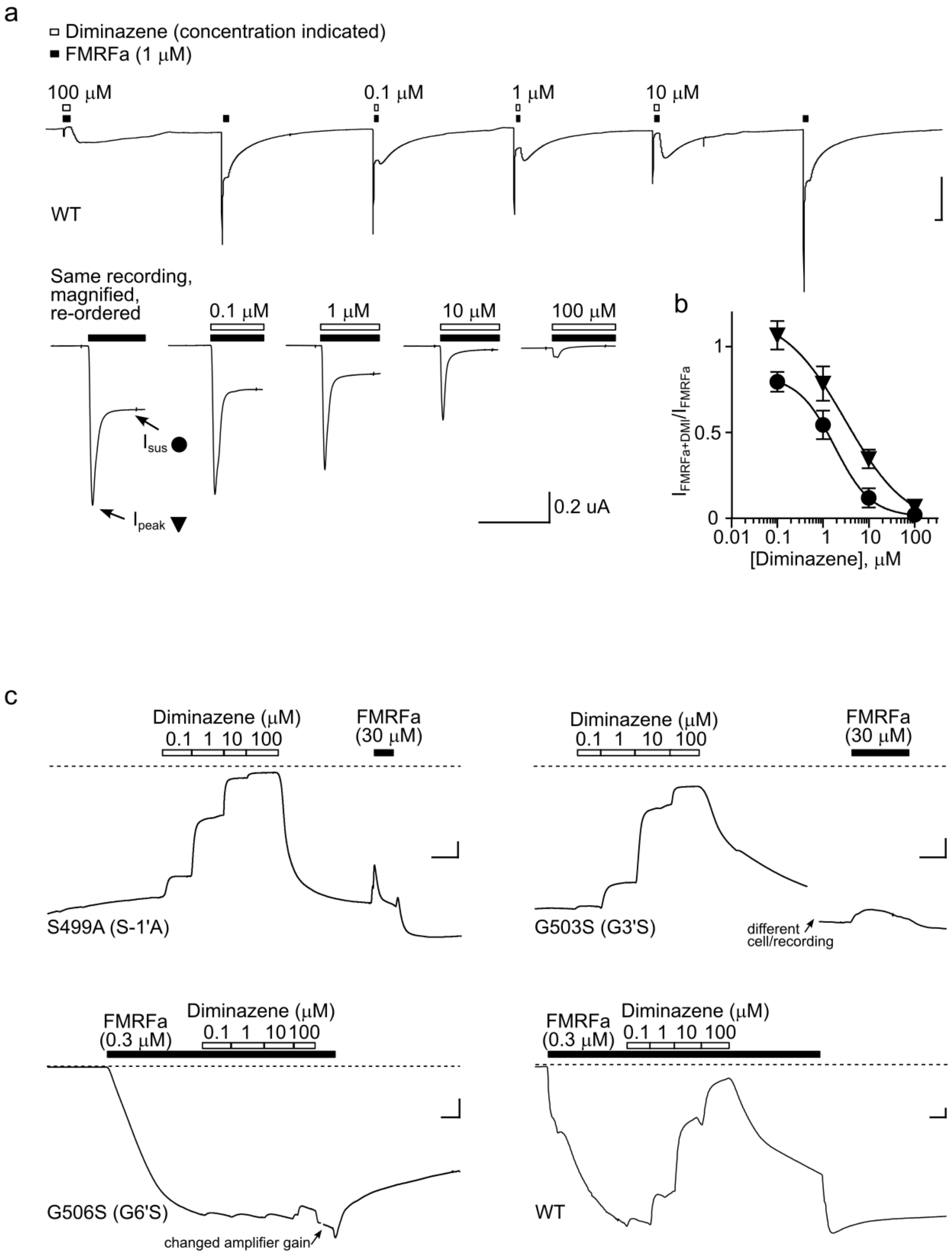


**Figure S6. Effects of diminazene on FaNaC1 channels**

(a) 1 μM FMRF-gated currents with and without diminazene in an oocyte expressing wildtype (WT) FaNaC1. Lower panel shows a magnified view, illustrating the inhibition of peak and sustained current by diminazene. (b) Mean ± SEM (n = 4) normalized peak and sustained current amplitude in the presence of increasing concentrations of diminazene. (c) Effect of diminazene and FMRFa on constitutive currents in oocytes expressing indicated mutant FaNaC1 channels (upper panel) and effect of diminazene on sustained 0.3 μM FMRFa-gated currents in oocytes expressing mutant or WT FaNaC1 (lower panel; mean ± SEM shown in Fig. 4d).

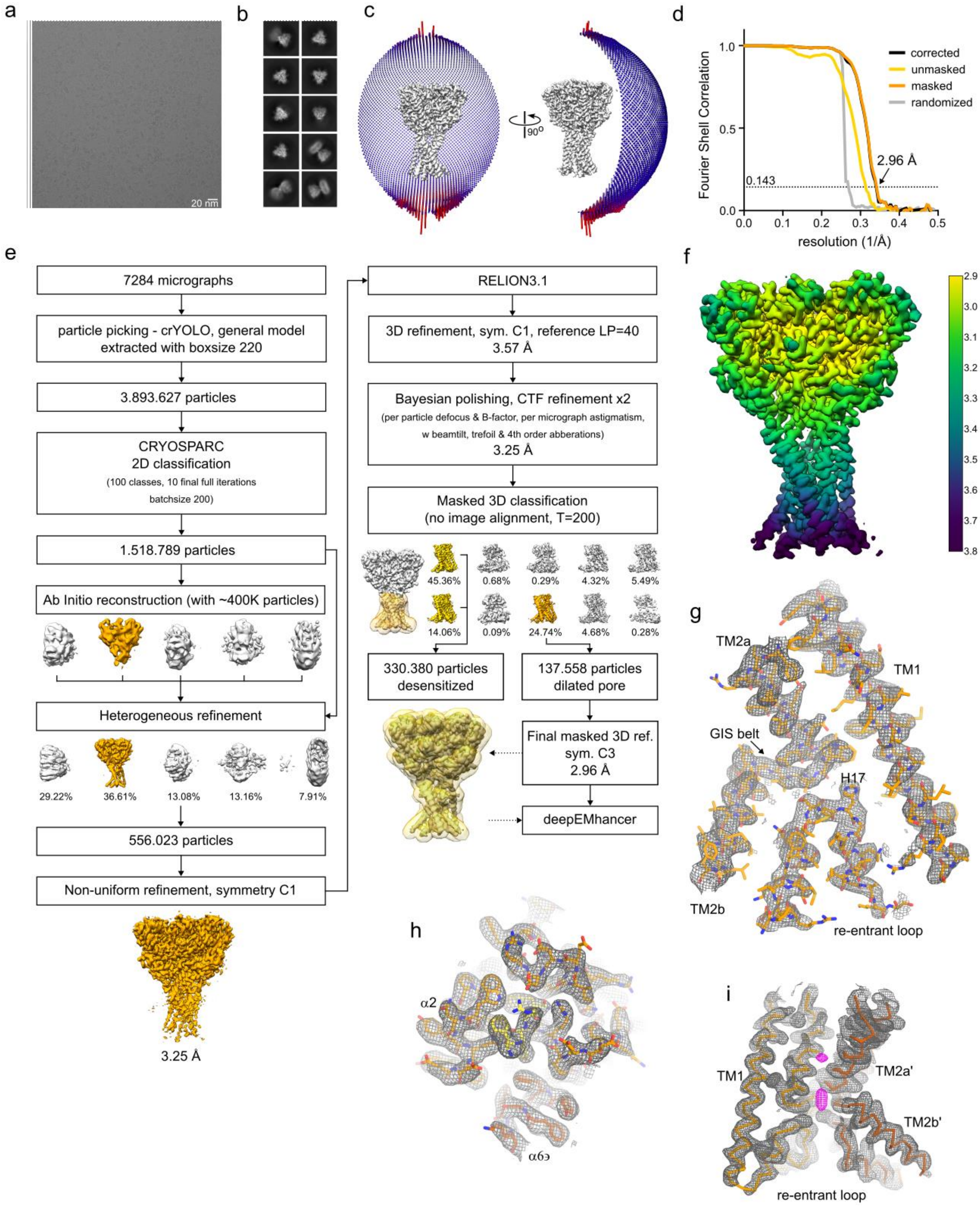


**Figure S7. FMRFa-bound FaNaC1 in the presence of diminazene, determination by cryo-EM**

Representative cryo-EM image (a) and 2D classes (b) of vitrified FaNaC1 in FMRFa-bound state supplemented with diminazene. (c) Angular distribution of the particles included in the final C3-symmetrised map. The length and the colour of the sticks represent the number of particles. (d) FSC plot used for resolution estimation (0.143 cut-off criteria). (e) Schematic representation of the processing workflow, the mask used in the final refinement iteration is displayed as a transparent outline. (f) Final deepEMhancer-postprocessed map coloured according to the local resolution estimation in Relion. (g) Density corresponding to the transmembrane domain shown as grey mesh, deepEMhancer map used for visualization is contoured at 4 $\sigma$. The respective fitted model is shown in orange. H. Close-up view of the ligand-binding site, deepEMhancer map is displayed as grey mesh contoured at 4 $\sigma$, fitted atomic model is displayed in orange, FMRFa molecule is shown in yellow. (i) Unassigned density in the pore region displayed in magenta, the surrounding protein density in grey, fitted atomic model is shown as C$\alpha$-trace in hues of orange. One subunit is not displayed for clarity.

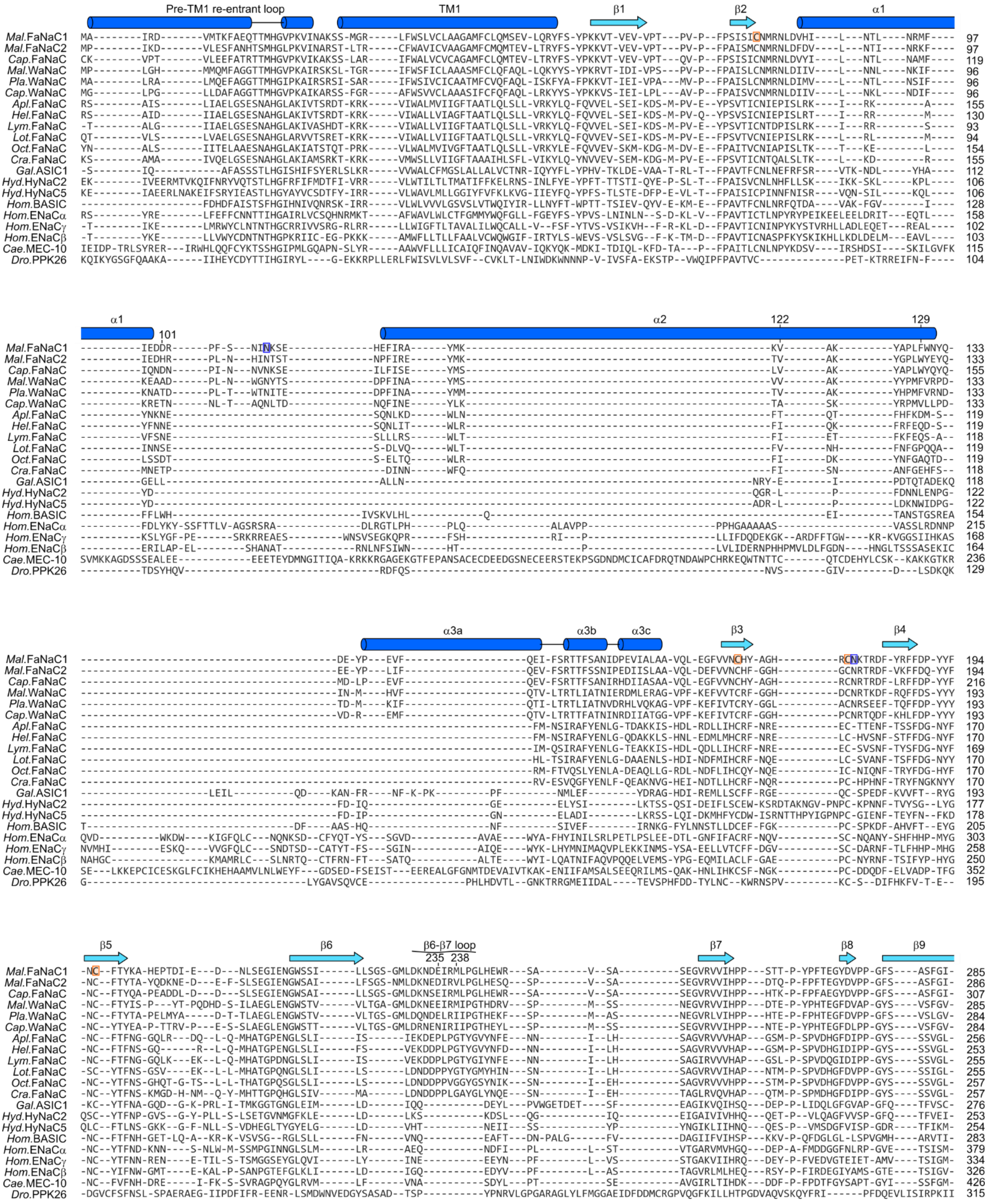


**Figure S8. (Continued overleaf)**

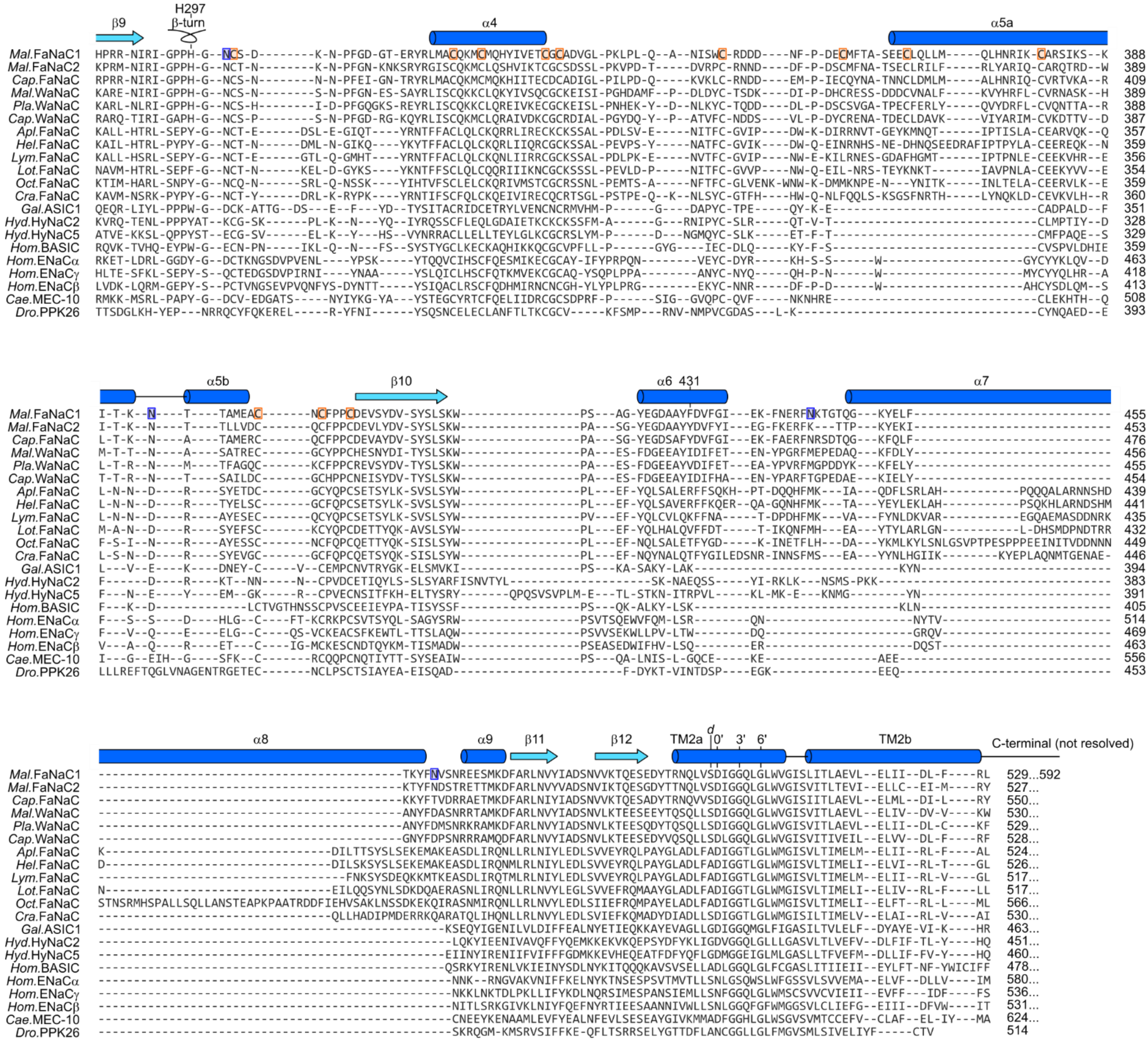


**Figure S8. Amino acid sequence alignment of FaNaCs and other DEG/ENaCs.** The following 21 DEG/ENaC sequences were extracted from an existing alignment of 544 diverse genes[1]. Six annelid genes from the FaNaC branch: *Malacoceros fuliginosus* (*Mal.*, bristle worm), *Capitella teleta* (*Cap.*, bristle worm), and *Platynereis dumerilii* (*Pla.*, clam worm) FMRFamide-gated $Na^+$ channels (FaNaCs) and Wamide-gated $Na^+$ channels (WaNaCs). Six mollusc genes from the FaNaC branch: *Aplysia kurodai* (*Apl.*, slug), *Helix aspersa* (*Hel.* snail), *Lymnaea stagnalis* (*Lym.*, snail), *Lottia gigantea* (*Lot.* limpet), *Octopus bimaculoides* (*Oct.*), and *Crassostrea gigantea* (*Cra.*, oyster) FaNaCs. Several diverse DEG/ENaC genes from various animals: *Gallus gallus* acid sensing ion channel 1 (*Gal.* ASIC1, chicken), *Hydra vulgaris* $Na^+$ channel subunits 2 and 5 (*Hyd.* HyNaC2,5), human (*Hom.*) bile acid-sensing ion channel (BASIC) and epithelial $Na^+$ channel (ENaC) subunits α, β, and γ, *Caenorhabditis elegans* mechano-sensitive channel 10 (*Cae.* MEC-10), and *Drosophila melanogaster* pickpocket 26 (*Dro.* PPK26). Secondary structure from our *Malacoceros* FaNaC1 structures is indicated above alignment. Certain FaNaC1 residues discussed in main text are indicated. TM2 prime numbering allows better comparison across channels[2]. *d* indicates position of gain-of-function *degenerin* mutations[3]. Orange boxes, cysteine residues forming disulfides. Blue boxes, glycosylation sites.

1 Dandamudi, M., Hausen, H. & Lynagh, T. Comparative analysis defines a broader FMRFamide-gated sodium channel family and determinants of neuropeptide sensitivity. *The Journal of biological chemistry* **298**, 102086, doi:10.1016/j.jbc.2022.102086 (2022).
2 Lynagh, T. *et al.* A selectivity filter at the intracellular end of the acid-sensing ion channel pore. *eLife* **6**, doi:10.7554/eLife.24630 (2017).
3 Eastwood, A. L. & Goodman, M. B. Insight into DEG/ENaC channel gating from genetics and structure. *Physiology (Bethesda)* **27**, 282-290, doi:10.1152/physiol.00006.2012 (2012).
